# Effects of greening-induced warming and cooling on tree phenology in temperate and boreal forests

**DOI:** 10.1101/2023.12.03.569807

**Authors:** Jing Guo, Jinmei Wang, Yuxin Qiao, Nicholas G. Smith, Zhiyong Liu, Rui Zhang, Xiuzhi Chen, Chaoyang Wu, Lei Chen

## Abstract

Tree phenology, periodic biological events in trees, is highly sensitive to climate change. It has been reported that forest greening can influence the local climate by altering the seasonal surface energy budget. However, tree phenological responses to forest greening remains poorly understood at large spatial scales. Combining remote-sensing derived phenological and leaf area indices since 2001, herein we show that forest greening led to earlier spring (−1.05 ± 0.17 d) and autumn phenology (−1.95 ± 0.14 d) in temperate and boreal forests. Our results show that forest greening in winter and spring decreased surface albedo and thus resulted in biophysical warming that caused earlier spring leaf phenology. In contrast, forest greening in summer and autumn triggered biophysical cooling by enhancing evapotranspiration, which led to earlier autumn leaf phenology. These findings suggest that forest greening could significantly alter tree phenology through seasonal biophysical impacts. Therefore, it is essential to incorporate these complicated biophysical impacts of greening into tree phenology models to accurately predict future shifts in tree phenology under future climate change.

## Introduction

Changes in tree phenology, periodical biological events in trees, affect not only growth and distribution of trees, but also biogeochemical processes of forest ecosystems, such as water, energy, and carbon cycling (1–3). Under global warming, shifts in tree phenology, such as earlier spring leaf-out or delayed leaf senescence, have been widely observed in temperate and boreal forests (4, 5). It is therefore critical to understand the climate-phenology relationship to accurately assess and predict the impacts of future climate change on forest ecosystems.

Under climate warming, numerous studies have reported vegetation greening and enhanced vegetation activities during the growing season, with a global and persistent increase of 8% in leaf area index (LAI) over the past 30 years (6–8). Warming-induced widespread greening may affect local climate through various biophysical feedbacks including radiative processes (i.e., albedo) and non-radiative processes (i.e., latent and sensible heat fluxes) (9–11). Vegetation greening could amplify, counteract, or even reverse the climate benefits of carbon sequestration through biophysical impacts (6, 7, 11). As the largest carbon reservoir of terrestrial ecosystems, the biophysical impacts due to changes in forest cover have received increasing attention over recent years (9–14). Forest greening generally reduces the surface albedo, leading to enhanced absorption of shortwave radiation (13, 15). Dissipation of this extra energy through processes such as evapotranspiration (ET) or heat convection can result in surface cooling (10, 11). Conversely, if the dissipation processes are limited, the excess energy contributes to surface warming (14). These intricate biophysical impacts also vary geographically, and seasonally (9–11). For example, forest greening-induced changes in biophysical properties in tropical regions has the potential to cause local cooling (12), while in temperate and boreal regions, it tends to lead to local warming (16). However, the temperate and boreal impacts vary seasonally, with greening often resulting in moderate biophysical warming in winter but cooling in summer (9). The dominant role of temperature in tree phenology has been widely documented (4, 17–19). Therefore, forest greening-induced local warming or cooling may also affect tree phenological events in spring and autumn. However, previous studies have mainly focused on the responses of tree phenology to anthropogenic warming, and little is known about the potential effect of forest greening on tree phenology. This knowledge gap reduces the reliability of predictions of tree phenology and carbon cycling under future climate change.

Combining remote sensing-derived phenological and leaf area indices between 2001 and 2021, here we examined the effect of forest greening on tree phenology in temperate and boreal forests. To this end, we first examined differences in tree phenology across different greening gradients in temperate and boreal forests between 2001 and 2021 using remote sensing datasets. To clarify the mechanisms of the greening-induced changes in tree phenology, we further examined the differences in local temperature, surface albedo, and evapotranspiration between high and low greening gradients. We hypothesized that seasonal greening-induced biophysical warming or cooling would alter tree phenology in temperate and boreal forests.

## Results

We calculated the mean annual differences in phenological indicators across different greening gradients in temperate and boreal forests from MODIS in 2001-2021, which represents potential shifts in tree phenology in response to greening induced by leaf area index (LAI) (Fig. 1). We found that compared with low greening areas, approximately 66.7% of high greening areas showed earlier start of the growing season (SOS), and about 33.3% of high greening areas showed delayed SOS (Fig. 1*A*). As with autumn phenology, we observed that approximately 81.6% of high greening areas showed advanced end of the growing season (EOS), and 18.4% of high greening areas had delayed EOS in comparison to low greening areas (Fig. 1*B*). On average, the EVI-based SOS began 1.05 d earlier, and EOS ended 1.95 d earlier within high greening areas compared to within low greening areas (Fig. 1, *C* and *F*). Moreover, we also analyzed the phenological differences between forests with high and low greening gradients across various latitudes, and climate zones in temperate and boreal forests. We observed a decrease in both ΔSOS and ΔEOS with increasing latitude (Fig. 1, *D* and *G*). Also, boreal regions exhibited significantly lower ΔSOS and ΔEOS compared to the temperate regions (Fig. 1, *E* and *H*). These results suggest a general phenomenon that the greening of temperate and boreal forests led to earlier spring and autumn phenology, and that the effects in boreal areas are greater than in other areas.

**Fig. 1.**
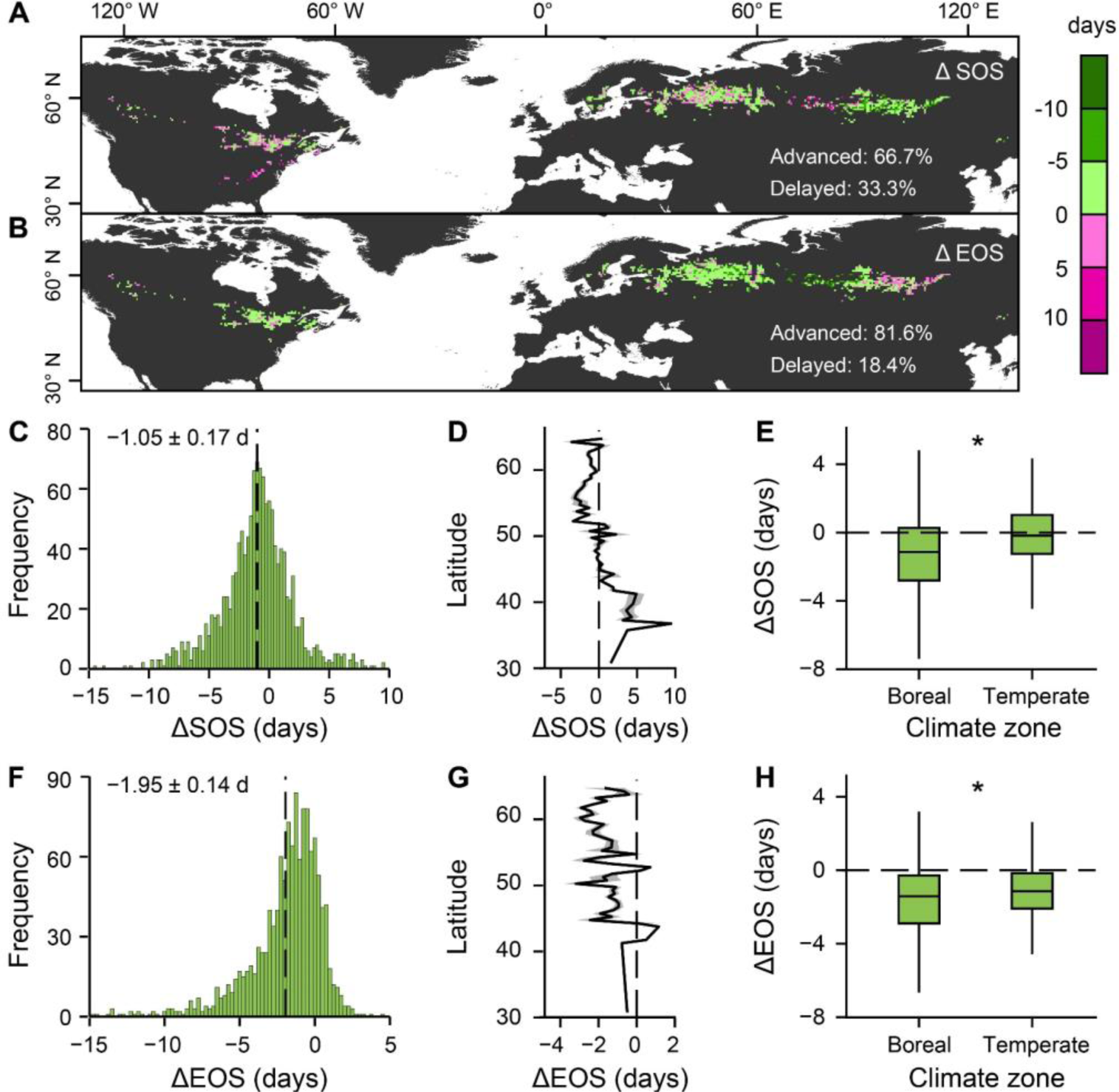
Spatial patterns and mean phenological differences across different greening gradients in temperate and boreal forests during the study period 2001-2021 across various latitudes, and climate zones. **A–H**, Spatial map of ΔSOS (**A**), and ΔEOS (**B**), distribution of ΔSOS (**C**), and ΔEOS (**F**), changes with latitude in ΔSOS (**D**), and ΔEOS (**G**), and changes in ΔSOS (**E**), and ΔEOS (**H**) for various climate zones. Positive ΔSOS, and ΔEOS represent delayed phenology, whereas negative ΔSOS, and ΔEOS indicate advanced phenology in **A–H**. The black dash lines represent mean annual phenological difference (ΔSOS, and ΔEOS) in **C** and **F**. Solid lines and shaded areas represent the mean and SD in **D** and **G**. The length of each box indicates the interquartile range, the horizontal line inside each box the median, and the bottom and top of the box the first and third quartiles respectively in **E** and **H**. The asterisk indicates a significant difference in the ΔSOS, and ΔEOS between temperate and boreal forests in **E** and **H** (*P*<0.01).

To clarify the underlying mechanisms of the greening-induced changes in tree phenology, we first examined the differences in seasonal LAI and daily land surface temperature (LST) between high and low greening gradients from MODIS in 2001-2021 (Fig. 2). The mean greening gradient during winter and spring (WS) in temperate and boreal forests is 0.23 m^2^/m^2^, which is lower than the value during summer and autumn (SA) (0.36 m^2^/m^2^; Fig. 2*E*). Meanwhile, we found that WS greening generated biophysical warming (0.76 ± 0.03°C), whereas greening in SA had weak biophysical cooling (−0.05 ± 0.01°C) (Fig. 2*H*). Spatially, we found that approximately 96.3% of high greening areas showed biophysical warming in WS, and about 62.8% of high greening areas had biophysical cooling in SA in comparison to low greening areas (Fig. 2, *C* and *D*). Using linear regression model, we also observed that the biophysical warming in WS significantly increased, while biophysical cooling in SA significantly decreased with the increase in greening gradient (Fig. 2, *F* and *I*). Furthermore, we used the near-surface air temperature (T_air_) to test the seasonal biophysical impacts of forest greening, and found similar results (Fig. S1). To shed light on the drivers of the seasonal biophysical impacts, we further examined the effects of seasonal greening on albedo and evapotranspiration. We found a significant decrease in albedo during the WS seasons in high greening areas compared to low greening areas (Fig. 2*J*). However, no significant difference in evapotranspiration was observed between high and low greening gradients (Fig. 2*G*). Furthermore, we found that evapotranspiration in the SA seasons showed a statistically significant increase with the increased greening, while no significant difference in albedo was observed between high and low greening gradients (Fig. 2, *G* and *J*). Therefore, the greening-induced biophysical warming in winter and spring is driven by decreased albedo (−0.07 ± 0.002), whereas the observed biophysical cooling is a result of increased evapotranspiration (0.44 ± 0.01 mm) in summer and autumn.

**Fig. 2.**
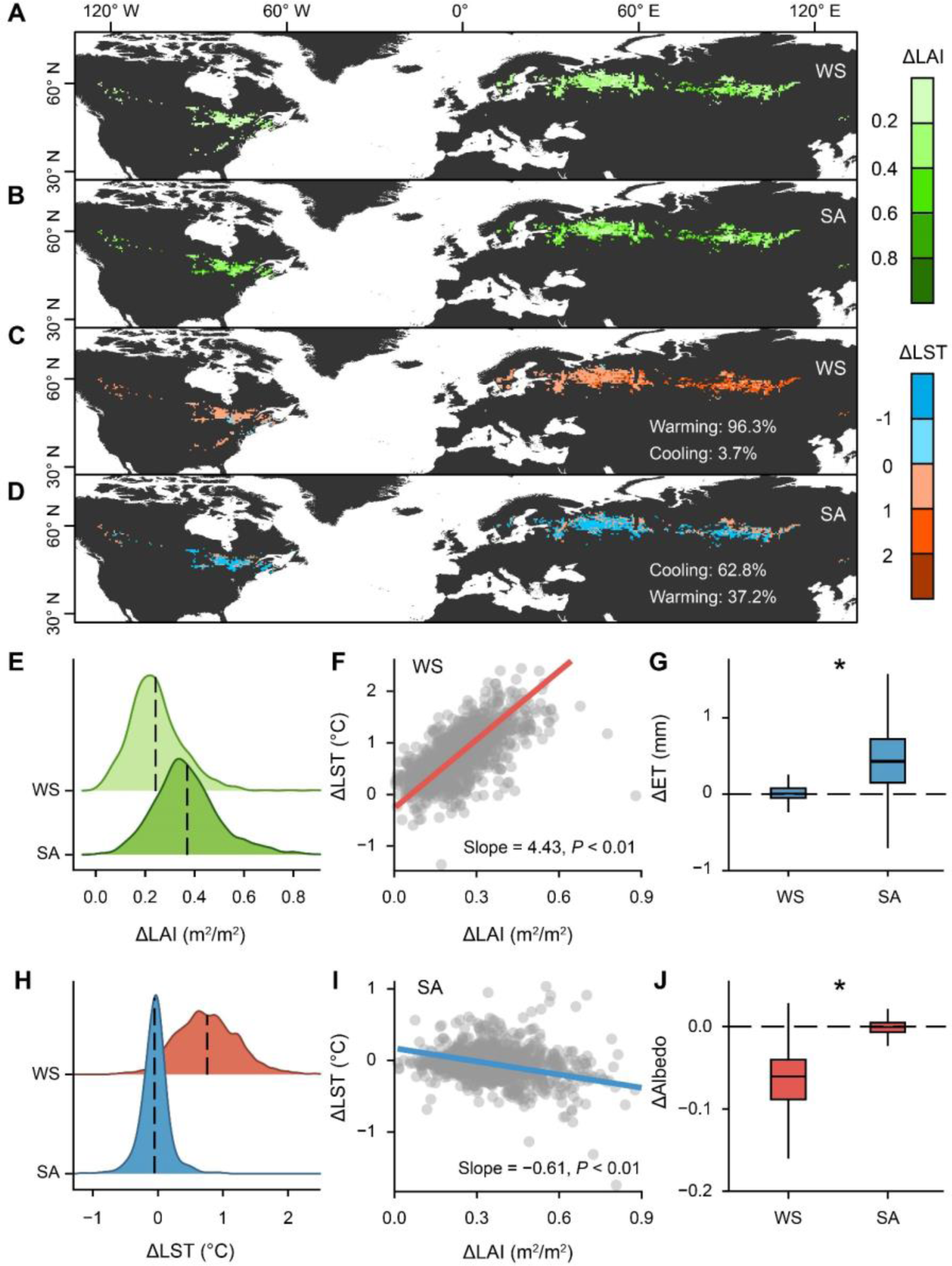
Effects of seasonal greening on biophysical impacts, evapotranspiration (ET), and albedo in temperate and boreal forests during the study period 2001-2021. **A**–**J**, Spatial map of average ΔLAI in winter and spring (WS, November to April) (**A**), and in summer and autumn (SA, May to October) (**B**), spatial map of average ΔLST in WS (**C**), and in SA (**D**), density plots of ΔLAI (**E**) and ΔLST (**H**) between WS and SA, changes in ΔLST with increase of **Δ**LAI in WS (**F**), and in SA (**I**), and changes in ΔET (**G**) and ΔAlbedo (**J**) between WS and SA. Positive ΔLST bothin WS and SA represents biophysical warming, whereas negative ΔLST in WS and SA indicate biophysical colling. The black dash lines in **E**, and **H** represent mean greening and temperature gradients for two growing seasons. In **F** and **I**, the circles represent the values of mean ΔLST in WS and in SA at each window. In **G** and **J**, the length of each box indicates the interquartile range, the horizontal line inside each box the median, and the bottom and top of the box the first and third quartiles respectively. The asterisk indicates a significant difference ΔET and ΔAlbedo between WS and SA (*P*<0.01). The black dash lines indicate when ΔET and ΔAlbedo are equal to zero.

Using linear regression models, we then test the effect of seasonal greening and local temperature on tree phenology between forests with high and low greening gradients. We observed that as both ΔLAI and ΔLST increased in WS, the ΔSOS significantly decreased (*P*<0.01; Fig. 3, *A* and *B*). As with autumn phenology, we found a significant decrease in ΔEOS with the increase in ΔLAI and decrease in ΔLST (*P*<0.01; Fig. 3, *C* and *D*). Compared to traditional vegetation indices such as enhanced vegetation index (EVI), the Near-infrared Reflectance of Vegetation (NIRv) index is more effective in isolating vegetation signals from background noise. In order to minimize the uncertainties caused by single phenological data source, we also applied the same analysis using the NIRv-derived phenology dataset, and found consistent results (Fig. S2). As phenological variations might also affect biophysical feedbacks, and thus local temperature in temperate and boreal regions, we further confirmed the causal relationship between phenological changes and biophysical impacts across different greening gradient (Fig. S3). We found that the magnitude of seasonal ΔLST was not determined by changes in spring and autumn phenology, but rather by the greening gradient in the corresponding preseason (Fig. S3). These suggested that the observed phenological differences across different greening gradients were due to seasonal biophysical impacts induced by forest greening.

**Fig. 3.**
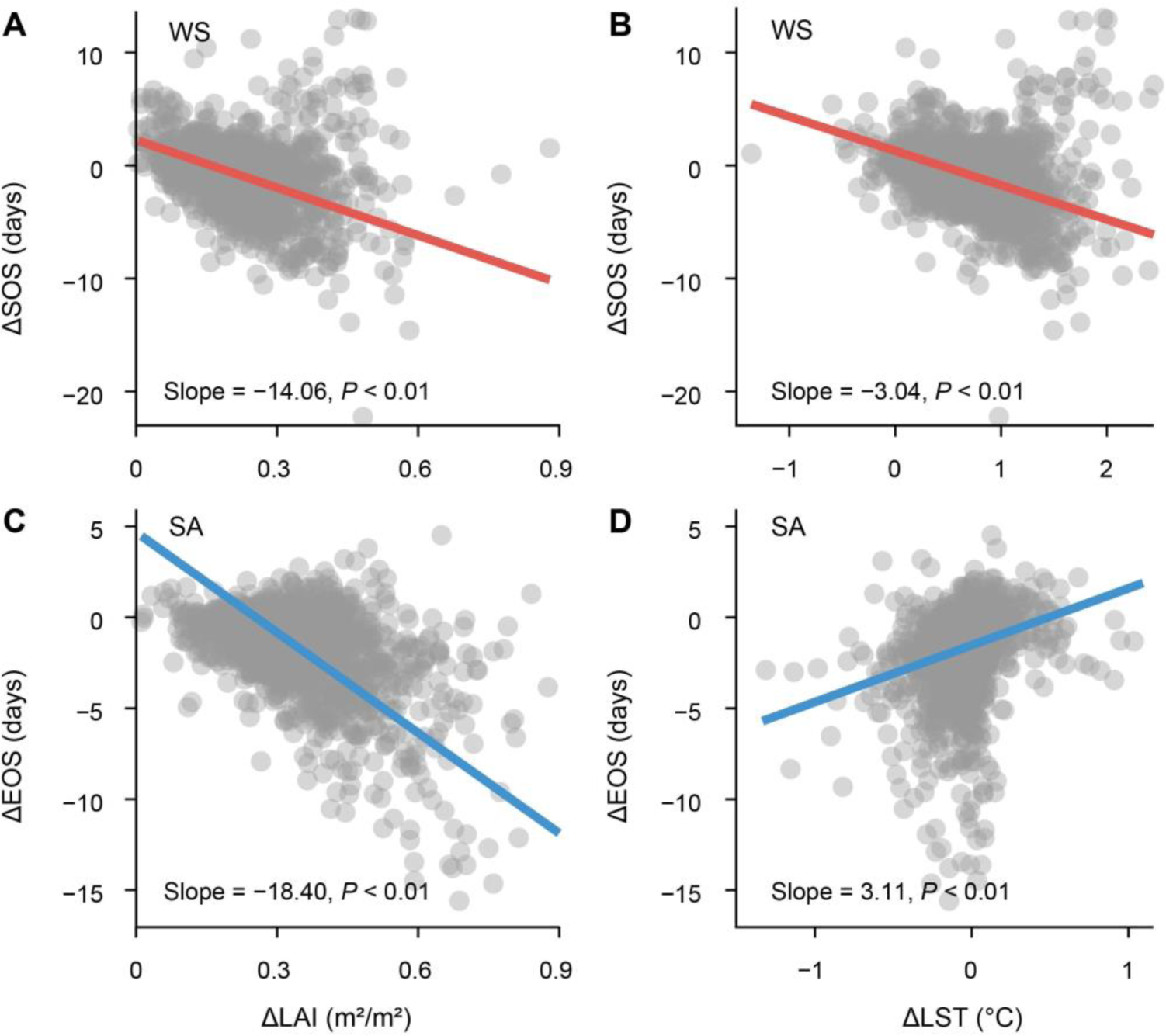
Effects of greening (leaf area index differences, ΔLAI) and greening-induced temperature gradient (land surface temperature differences, ΔLST) for two growing seasons on phenological differences across different greening gradients in temperate and boreal forests over the study period 2001-2021. **A–D**, Changes in ΔSOS with the increase in ΔLAI (**A**) and ΔLST (**B**) in winter and spring (WS, November to April), changes in ΔEOS with the increase in ΔLAI (**C**) and ΔLST (**D**) in winter and spring (SA, May to October). In **A** to **D**, the circles represent the values of mean ΔSOS in WS and ΔEOS in SA at each window.

Commonly, the time of spring leaf-out is primarily controlled by winter or spring temperature, while leaf senescence date is significantly influenced by summer or autumn temperature. We further examined the effect of greening on tree phenology between forests with high and low greening gradients for single seasons (Figs. S4, S5, S6). Similarly, we found a significant increase in ΔLST in response to greening in both winter and spring, with the highest increase observed in the spring season (Fig. S4, *A* and *B*). We also observed a significant decrease in ΔSOS when ΔLAI or ΔLST increased during both winter and spring season (Fig. S5). In particular, there was a more pronounced increase in ΔSOS in spring (Fig. S5, *C* and *D*). Additionally, we further observed that with an increase in the greening gradient, there was a greater decrease in ΔLST during summer compared to autumn (Figs. S4, *C* and *D*). The ΔEOS showed a significant decrease with an increase in ΔLAI and a decrease in ΔLST, particularly characterized by a notable decrease in ΔEOS during summer (Fig. S6). Our analyses again confirmed that WS greening, especially spring greening, could lead to an earlier SOS through biophysical warming. In contrast, the advanced EOS was attributed to greening-induced biophysical cooling in SA, with a more pronounced impact in summer.

**Fig. 4.**
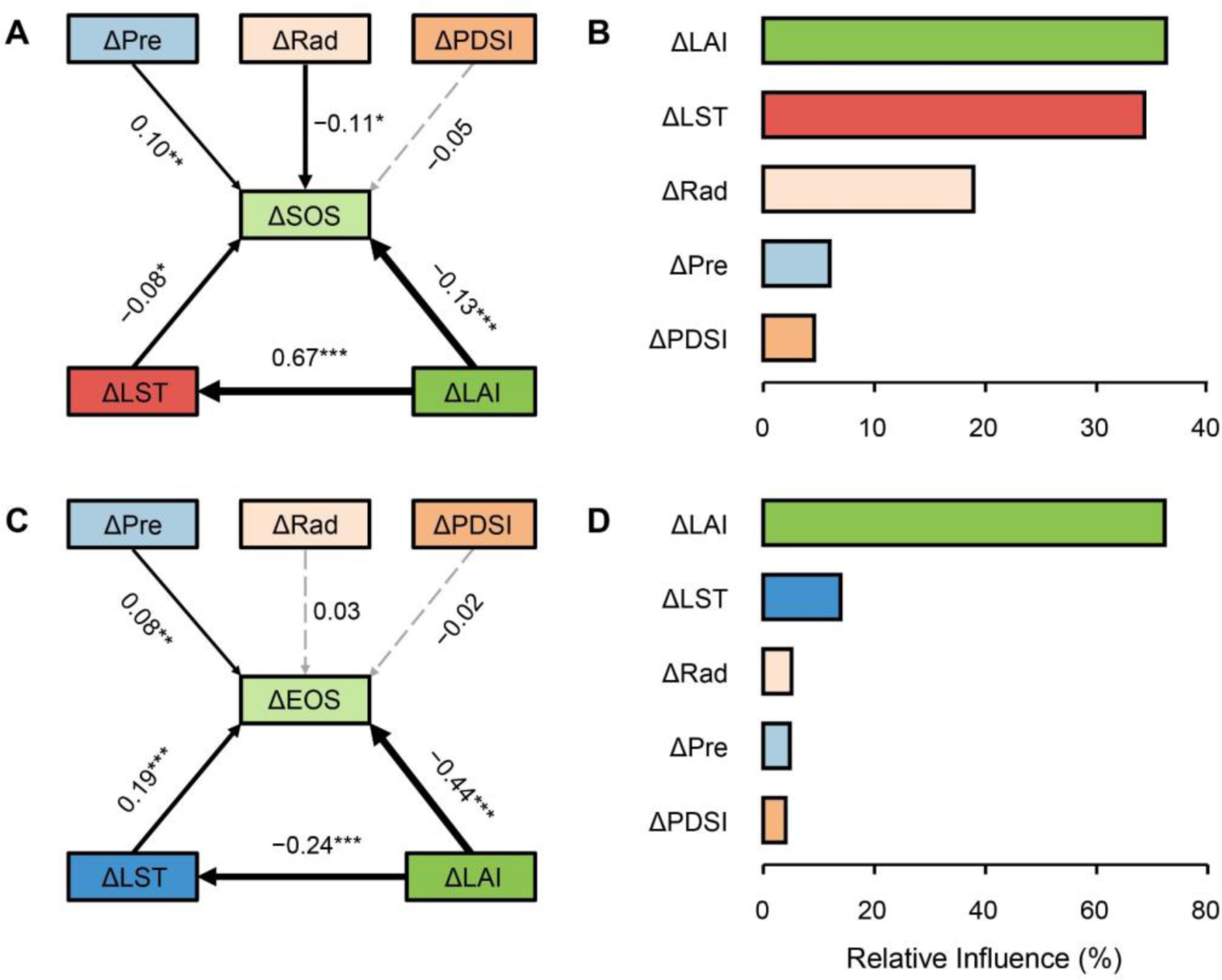
Effects of forest greening and climate variables gradients on phenological differences across different greening gradients in temperate and boreal forests during the study period 2001-2021. **A–D**, Piecewise structural equation model (SEM) for ΔSOS (**A**) and ΔEOS (**C**) considering both forest greening (ΔLAI) and climate variables gradients, relative influence of forest greening (ΔLAI) and climatic factors during previous growing on ΔSOS (**B**) and ΔEOS (**D**). In **A** and **C**, both ΔLAI and climate variables gradients (ΔLST, ΔPre, ΔPDSI, ΔRad) were incorporated into the SEM to test the direct (arrows from each climate factor gradient directly point to the ΔSOS or ΔEOS) or indirect (arrows from ΔLAI firstly directly point to ΔLST then to the ΔSOS or ΔEOS) effects of forest greening, and other climate factors gradients in fixed season on ΔSOS (**A**), and ΔEOS (**C**), with green lines indicating a negative effect and orange lines indicating a positive effect. The solid lines represent significant relationships (*P* < 0.05) between variables, while dashed lines represent no significant relationships between variables (*P* > 0.05). The calculated *P* values based on two-sided test and other statistics were listed in Table S1 and S2. In **B** and **D**, boosted regression trees (BRT) was used to quantify and compare the effects of climate variables gradients and forest greening on ΔSOS (**B**) or ΔEOS (**D**).

**Fig. 5.**
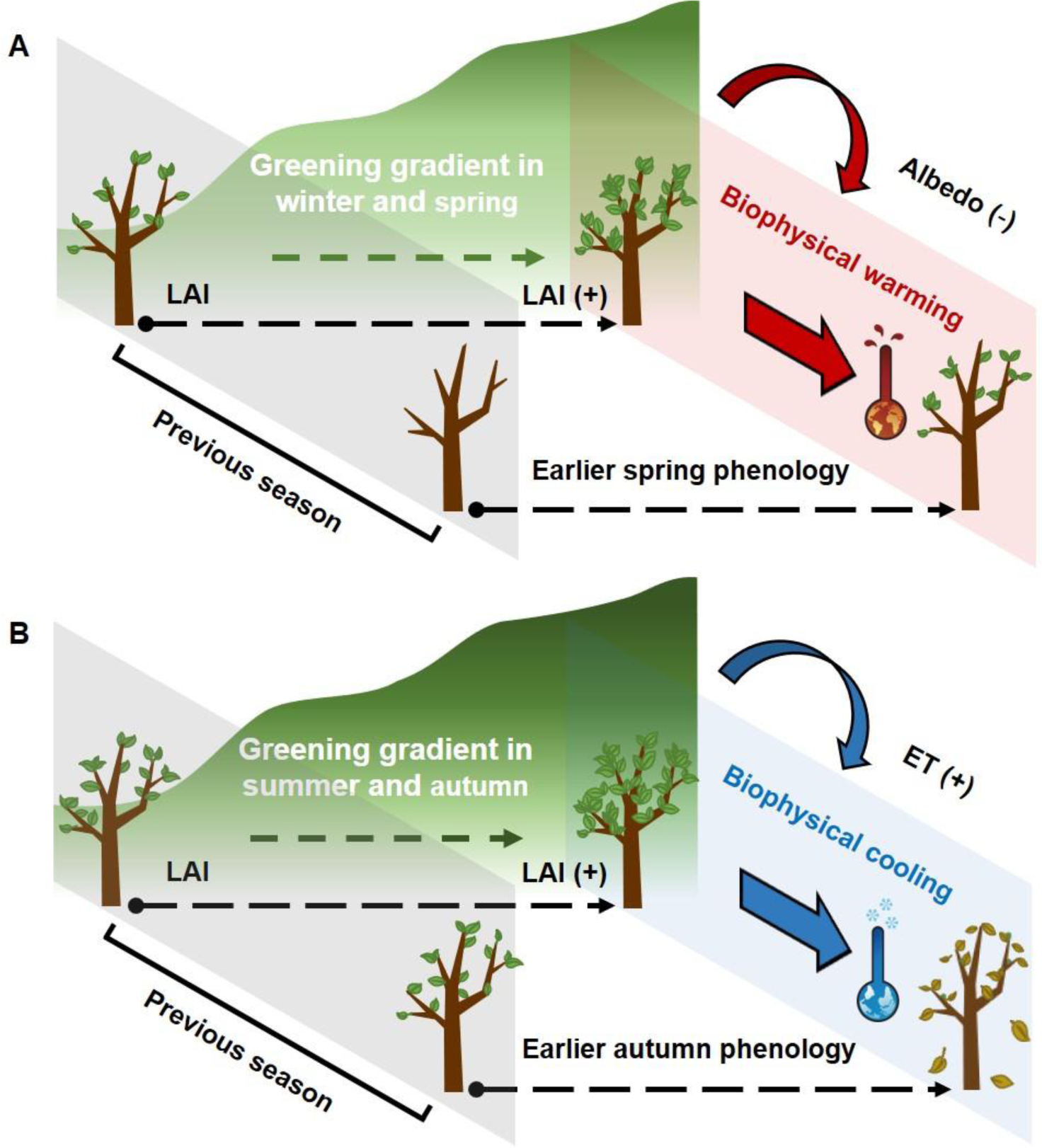
A schematic diagram of tree phenology in response to greening-induced biophysical impacts of previous season. **A–B**, Greening and biophysical warming in winter and spring drivers earlier spring phenology (**A**), greening and biophysical cooling in summer and autumn lead to advanced autumn phenology (**B**).

Phenological differences across different greening gradients are probably driven from other climate factors. To ensure the robustness of our results, we constructed a piecewise structural equation model (SEM) to investigate the direct and indirect effects of forest greening and climate factors gradients on ΔSOS and ΔEOS (Fig. 4, *A* and *C*). We observed that SOS was advanced directly by ΔLAI, ΔLST and ΔRad, but delayed by ΔPre in the WS seasons. The direct effect of ΔPDSI on SOS was not significant (Fig. 4*A*). We again found that increased LAI significantly increased LST. SOS was significantly advanced by increased LST, which provides robust evidence for indirect effect of ΔLAI on SOS through biophysical warming. Furthermore, we found that EOS was advanced directly by ΔLAI, and ΔLST, but delayed by ΔPre in the preseason (SA). The direct effects of ΔPDSI and ΔRad on EOS were not significant (Fig. 4*C*). We similarly found the indirect effect of LAI through biophysical cooling. The increased LAI significantly decreased LST, and thus advanced EOS. Using boosted regression tree (BRT) models, we further analyzed and compared the relative importance of ΔLAI, ΔLST, and other climatic gradients to ΔSOS and ΔEOS, respectively. Results showed that both ΔSOS and ΔEOS were mainly attributed to seasonal ΔLAI and ΔLST (Fig. 4, *B* and *D*).

## Discussion

Over recent decades, warming-induced shifts in tree phenology have been widely observed in temperate and boreal forests (4, 5, 17). However, previous studies focused mainly on responses of tree phenology to anthropogenic warming, but have failed to explore the effect of greening-induced warming or cooling on tree phenology. Using long-term and large-scale phenological and leaf area datasets, herein we demonstrated that forest greening significantly advanced start of the growing season (SOS) and end of the growing season (EOS) through seasonal biophysical warming and cooling, respectively, in temperate and boreal forests (Fig. 5).

In temperate and boreal forests, earlier spring phenological events, such as leafing or flowering, due to anthropogenic warming has been observed across multiple taxa and regions (5, 17, 18). This is because warming can accelerate the accumulation of thermal units required to break ecodormancy, and thus cause an earlier spring phenology (5, 20, 21). Here we observed that SOS occurred earlier with the increased greening in winter and spring. A related idea that might explain the greening-induced earlier spring phenology is the biophysical warming due to forest greening. To test this hypothesis, we calculated and compared the difference in land surface temperature in winter and spring seasons between forests with high and low greening gradients. We found local temperature showed a significant increase with the increased greening. This suggested that greening-induced biophysical warming led to the observed earlier spring phenology. This effect was supported by the observed negative correlation between greening-induced land surface temperature and dates of SOS. Generally, forest greening can affect local temperature by altering surface biophysical properties (i.e., albedo and evapotranspiration) (7, 9). To shed light on the drivers of the biophysical warming, we further examined the effects of greening in winter and spring on albedo and evapotranspiration. We found a statistically significant reduction in albedo within high greening areas in comparison to low greening areas. However, no significant discrepancy in evapotranspiration was observed between high and low greening areas. This finding suggests that forest greening in winter and spring reduced surface albedo, and thus resulted in earlier spring phenology. We further analyzed the biophysical impacts of winter and spring greening, separately, and observed stronger effects of spring greening on local temperature and spring phenology than winter greening. This could be related to the widespread leaf-out in spring (5, 17, 20).

In addition, we observed that EOS occurred earlier with the increased greening in summer and autumn seasons. To elucidate the greening effect on EOS, we further examined the effect of greening in summer and autumn seasons on land surface temperature. We found forest greening in summer and autumn seasons, especially in summer, significant reduced local temperature. This greening-induced local cooling may accelerate the rate of chlorophyll degradation, reduce the activities of photosynthetic enzymes, and increase the risk of late autumn frost, ultimately advancing autumn phenology (4, 22–24). This was also supported by the observed positive correlation between greening-induced land surface temperature and dates of EOS. Furthermore, we found evapotranspiration showed a significant increase with the increased greening, while no significant greening effect on albedo was observed. These findings suggested that greening-induced increases in evapotranspiration in summer and autumn triggered the biophysical cooling, and thus led to the observed earlier autumn phenology within high greening areas compared to within low greening areas. Nevertheless, we observed that summer greening had stronger biophysical impacts than autumn greening. Correspondingly, effect of biophysical cooling in summer on EOS was also stronger than that in autumn season. This could be related to the higher greening and enhanced evapotranspiration in summer than that in autumn (Fig. S7).

Moreover, we also observed a greater advance in both spring and autumn phenology in boreal zones compared to temperate zones. We calculated greening and land surface temperature differences between forests with high and low greening gradients in boreal and temperate areas, respectively (Fig. S8). We found that forest greening in winter and spring and greening-induced biophysical warming in boreal areas were significantly higher than in temperate areas (Fig. S8, *A* and *B*). This results in greater effects on spring phenology in boreal areas compared to temperate areas. Additionally, we found that both summer and autumn greening and corresponding cooling were significantly lower in boreal zones than in temperate zones (Fig. S8, *C* and *D*). However, the phenology of species in colder regions is likely to be more sensitive to temperature variation than in warmer regions (25–27). Therefore, the lower extent of greening-induced biophysical cooling in boreal forests may cause an earlier autumn phenology compared to temperate forests.

Temperature has long been recognized as the primary environmental cue for tree phenology (17–19). However, in addition to temperature, shifts in tree phenology across different greening gradients are probably influenced by other climate factors. To further test the greening-driven hypothesis, we constructed an LAI-based SEM model to examine the relationships between greening gradient, gradients in climate factors, and phenological differences. As previously mentioned, we similarly found that the seasonal greening has a direct effect on both spring and autumn phenology, but also an indirect effect through the land surface temperature. We compared the relative importance of greening and climate factors to phenological differences. These results further emphasized the importance of greening in tree phenology. These findings suggested that forest greening could significantly alter tree phenology through seasonal biophysical warming and cooling. In recent decades, the process-based phenological models has greatly improved our ability to predict phenological shifts in response to climate warming (4, 19, 25, 28). However, these models often based on temperature changes due to anthropogenic warming (25), but neglected the seasonal biophysical warming and cooling induced by greening. Hence, the interactions between tree phenology and biophysical impacts needed to be well represented in tree phenology models to better predict the future shifts in tree phenology in a warmer world.

Combining remotely sensed phenological and leaf area indices between 2001 and 2021, we found that forest greening led to earlier spring and autumn phenology in temperate and boreal forests. The earlier spring phenology was driven by forest greening-induced reductions in winter and spring surface albedo that caused biophysical warming. In contrast, summer and spring forest greening induced biophysical cooling by increasing evapotranspiration, which led to earlier autumn phenology. Our results demonstrate that forest greening could significantly alter tree phenology through seasonal biophysical impacts. Moreover, our findings emphasize the crucial role of leaf area index as a key predictor in understanding the changes in tree phenology in a warmer world. It is therefore essential to incorporate these complicated biophysical impacts of greening into tree phenology models to accurately predict future shifts in tree phenology under future climate warming scenarios.

## Materials and Methods

### Land cover type product

The MODIS land cover product (MCD12Q1) with IGBP classification at 500m spatial resolution was used to distinguish the forested regions (29). According to the land cover map in 2001, we excluded the pixels representing urban lands, grasslands, crops, and water bodies. This left five forest types: evergreen needleleaf forests (ENF), evergreen broadleaf forests (EBF), deciduous needleleaf forests (DNF), deciduous broadleaf forests (DBF), and mixed forests (MF) (Fig. S9).

### Leaf area index dataset

Leaf area index (LAI) has been widely used to characterize the vegetation greenness (8). LAI data in this study between 2001 and 2021 were obtained from Terra and Aqua MODIS LAI products (MOD15A2H and MYD15A2H) at 500 m spatial resolution and 8-day temporal resolution (30). We used quality assurance (QA) flag of LAI products to remove low quality data contaminated by clouds, aerosols, shadows, and snow.

### Phenology dataset

Our phenology dataset was the land surface phenology from MODIS Land Cover Dynamics products (MCD12Q2), with a spatial resolution of 500 m between 2001 and 2021 on a global scale (31). The phenological metrics were derived from the 8-day Enhanced Vegetation Index (EVI), which is calculated using MODIS Nadir Bidirectional Reflectance Distribution Function (BRDF) adjusted surface reflectance (NBAR-EVI2). The penalized cubic smoothing splines were used to fit the 8-day MODIS-EVI time series and extract the start of growing season (SOS) and end of growing season (EOS). The SOS was defined as the date when the fitted EVI2 time series first crossed 15% of the segment EVI2 amplitude, and EOS was defined as the date when the fitted EVI2 time series last crossed 15% of the segment EVI2 amplitude.

To reduce the uncertainties resulting from a single data source, we also extracted phenological metrics using Near-infrared Reflectance of Vegetation (NIRv) dataset. The NIRv was a newly developed vegetation index, which is more sensitive to distinguish vegetation signals from background noise (32). Compared to traditional NDVI and EVI datasets, the NIRv dataset showed a higher accuracy in phenology estimation (33). NIRv data in this study between 2001 and 2021 were derived from MODIS Nadir BRDF-Adjusted Reflectance (NBAR) products (MCD43A4) with 500 m spatial resolution and daily temporal resolution (34). We used quality assurance (QA) flag to exclude the effect of atmosphere on the data. The NIRv was calculated as below:

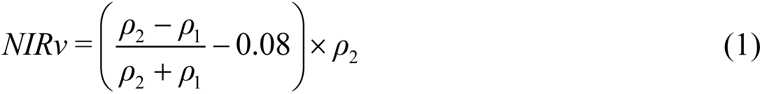

where *ρ*_1_ and *ρ*_2_ represent surface reflectance of MODIS band 1 (620-670 nm) and 2 (841–876 nm), respectively (35).

We used Savizky-Golay smooth method (36) to minimize the noise of atmospheric interference and satellite sensor before the estimation of spring and autumn phenology. We applied a double logistic function (Eq. 2) to fit time series NIRv, and then extracted SOS and EOS (37). The SOS is defined as the timing of first local maximum point in the first half year, and the date of second local maximum point in the second half year is defined as the EOS.

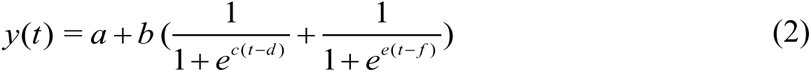

where *b*, *c*, *d*, and *f* are parameters of logistic function, *a* represents the initial background NIRv value, *a + b* denotes the maximum NIRv value, *t* is time in days, and *y(t)* is the NIRv value at time *t*.

### Climate zones

The Global ecological zone (GEZ) map at a 1 km spatial resolution was used to define the climate zones (38). We only kept extra-tropical regions (i.e., latitudes >30°N), which is characterized by distinct seasonal phenological cycles. We reclassified forest biomes into subtropical forests, temperate forests, and boreal forests. Given the limited number of screened windows available in subtropical regions, we focused on temperate and boreal forests in the Northern Hemisphere (Fig. S10).

### Climate data

Daytime and nighttime land surface temperature (LST) data were derived from Terra and Aqua MODIS products (MOD11A1 and MYD11A1) with 1 km spatial resolution and 8-day interval from 2001 to 2021 (39). Daily LST was obtained as an average of daytime and nighttime LST with an error <1 K.

Evapotranspiration (ET) data were from MODIS ET products (MOD16A2) at 500 m spatial resolution and 8-day temporal resolution between 2001 and 2021, which was generated using air temperature, air pressure, air humidity, LAI, albedo, and land cover (40).

The MODIS albedo products (MCD43A3) provided black sky albedo and white sky albedo over shortwave broadband, with 500 m spatial resolution and 16-day interval from 2001 to 2021 (41). Because of the similarity between black– and white-sky albedo, we used the average of black– and white-sky albedo to represent actual albedo (9, 10).

Monthly near-surface air minimum and maximum temperatures (T_min_ and T_max_), shortwave radiation (Rad), precipitation (Pre), and Palmer Drought Severity Index (PDSI) between 2001 and 2021 were derived from TerraClimate datasets, with a high-spatial resolution of 4 km (42). The T_air_ was calculated as an average of T_min_ and T_max._

Digital elevation models (DEM) data were obtained from the GTOPO30 dataset with a 1 km spatial resolution (43). To reduce differences in the spatial resolutions between various remote sensing datasets, all satellite data were resampled to a spatial resolution of 4 km.

### Window searching approach

Window searching approach was applied to examine all available samples, and to compare high greening areas with low greening areas in temperate and boreal forests. The purpose of this search strategy was to exclude the differences in the climatic background between forests with high and low greening gradients (9). Following from previous studies (9, 10), the search window size is defined as 0.5° × 0.5° (longitude and latitude, respectively). We then screened all windows according to the following criteria: (1) we used land cover map (MCD12Q1) in 2001 to filter the windows that only contain forest pixels (i.e., ENF, EBF, DNF, DBF, and MF); (2) these selected windows had at least 70% fractional forest cover; (3) in order to minimize potential systematic bias in land surface temperature assessments, we constrained the elevation difference within 500 m in each window. To ensure the robustness of results, we also applied the same analysis using search windows with at least 30% and 50% fractional forest cover, respectively, and obtained similar results. Thus, we selected search windows with at least 70% fractional forest cover as representatives for subsequent analyses.

### Statistical analysis

We aggregated the 8-day composite LST, and LAI data to monthly mean values. To delineate the forest boundary of high and low greening areas within each screened window, we first calculated the mean annual LAI of forest within each screened window between 2001 and 2021 (Window_mean-LAI_). We defined the high greening areas as the region where LAI pixel values exceeded Window_mean-LAI_ within each screened window. Conversely, the areas with LAI pixel values lower than Window_mean-LAI_ were identified as low greening areas. We then established the monthly dynamic boundaries for the high and low greening areas in each screened window. The average phenological metrics (SOS, and EOS), and climate factors (LST, T_air_, Pre, PDSI, Rad, ET, Albedo, and DEM) within high and low greening areas were extracted according to the monthly forest boundary, respectively.

In temperate and boreal forests, previous November 1st is often used as the starting date of the preseason, a period during which temperature is related to spring leaf-out (17, 18). Across the forest pixels in all selected windows, the mean date of SOS was DOY 117. Therefore, the period between the previous November 1 and April 30 (winter and spring, WS) was considered as the preseason of spring phenology. Furthermore, it has been reported that the temperatures during summer and autumn have an impact on leaf senescence (44–46). The mean EOS across all selected windows was DOY 285 for the forest pixels. We used the period from May 1 to October 31 (summer and autumn, SA) as the preseason of autumn phenology. We screened the windows for spring and autumn phenology separately, according to the corresponding fixed preseason. Moreover, we also removed the records of SOS and EOS exceeding 2.5 times of median absolute deviation (MAD) to exclude potential biases caused by outliers, respectively (32). In total, 1344 windows were retained for spring phenology analysis, while 1274 windows were used for EOS analysis.

The space-for-time approach has been widely applied to examine the effect of land use/cover change and earth surface greening on local temperature (9, 10, 14). We used the “space-for-time” method to calculate mean phenological differences (ΔSOS and ΔEOS) between high and low greening areas within each window during 2001-2021 according to:

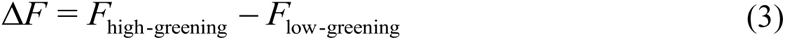

where Δ*F* represents the gradients between high and low greening areas for tree phenology (SOS and EOS), forest greenness (LAI), and climate factor (LST, Pre, PDSI, Pre, Rad, ET, and Albedo). *F*_high-greening_ represents these values in high greening areas, and *F*_low-greening_ represents these values in low greening areas. The changes in ΔSOS and ΔEOS were then analyzed across latitudes, and climate zones. One-way analysis of variance (ANOVA) was used to test the difference in ΔSOS and ΔEOS between temperate and boreal forests.

To clarify the underlying mechanisms of the greening-induced shifts in tree phenology, we examined the differences in seasonal LAI, and LST between high and low greening gradients. Specifically, we first calculated ΔLAI and ΔLST in the WS seasons (corresponding to SOS) and in the SA seasons (corresponding to EOS) within each window during 2001-2021 according to Equation (3). The changes in ΔLAI and ΔLST in different preseason were also analyzed across climate zones. One-way ANOVA was used to examine the difference in ΔLAI, and ΔLST between temperate and boreal forests. Using linear regression models, we further examined the relationships between seasonal greening (ΔLAI) and corresponding biophysical impacts (ΔLST). We also used the near-surface air temperature (T_air_) indicator to examine the seasonal biophysical impacts of forest greening, and obtained similar results. Due to the availability of fine-scale land surface temperature data, we used satellite-derived LST for subsequent analyses. To shed light on the drivers of the seasonal biophysical impacts, we calculated seasonal Δalbedo and ΔET between high and low greening gradients according to Equation (3). One-way ANOVA was used to test the difference in Δalbedo, and ΔET between the WS and SA seasons.

We further investigated the effects of seasonal greening and biophysical impacts on tree phenology. Linear regression model was used to test the relationship between shifts in phenology and seasonal greening (ΔLAI) and local temperature (ΔLST). Moreover, we analyzed the effect of greening on tree phenology shifts for single seasons (i.e., winter, spring, summer, and autumn) using linear regression models. To minimize the uncertainties rising from a single phenological data source, we used the same analysis to investigate the effect of greening on tree phenology using the NIRv phenological dataset. Given the potential impact of phenological variations on local temperature in temperate and boreal regions, we also examine the causal relationship between phenological changes and biophysical impacts across different greening gradients.

Phenological differences between forests with high and low greening gradients are probably driven from other climate factors. To test our greening-driven hypothesis, we used piecewise structural equation models (SEM) to further examine the direct and indirect effects of greening gradient, and other climate factors gradients in fixed preseason on tree phenology. We selected Rad, Pre, and PDSI as other controlling drivers of tree phenology. In the SEM model, we hypothesized that seasonal greening and climate variables are likely to directly influence SOS and EOS, indicated by the arrows from ΔLAI, ΔLST, ΔRad, ΔPre, and Δ PDSI directly point to the ΔSOS or ΔEOS. Also, forest greening can indirectly influence SOS and EOS by altering the seasonal biophysical impacts, indicated by the arrows from ΔLAI firstly directly point to ΔLST, then to the ΔSOS and ΔEOS. The piecewise SEM was conducted using the “piecewiseSEM” package (47) in R (48). Further, we quantified and ranked the effects of seasonal greening and these climate factors on tree phenology using boosted regression trees (BRT), an ensemble learning method that incorporate both statistical and machine learning techniques (49). We conducted BRT analysis using “gbm” package (50) in R (48).

All the data analyses were conducted using Google Earth Engine (51) and R version 4.1.2 (48).

## Author contributions

L.C. designed the research. J.G., J.W. and Y.Q. performed the data analysis. J.G. wrote the paper with the inputs of J.W., Y.Q., N.G.S., Z.L., R.Z., X.C., C.W. and L.C. All authors contributed to the interpretation of the results and approved the final manuscript.

## Acknowledgments

This work was funded by National Natural Science Foundation of China (32271833).

## Supplementary Information

**Fig. S1.**
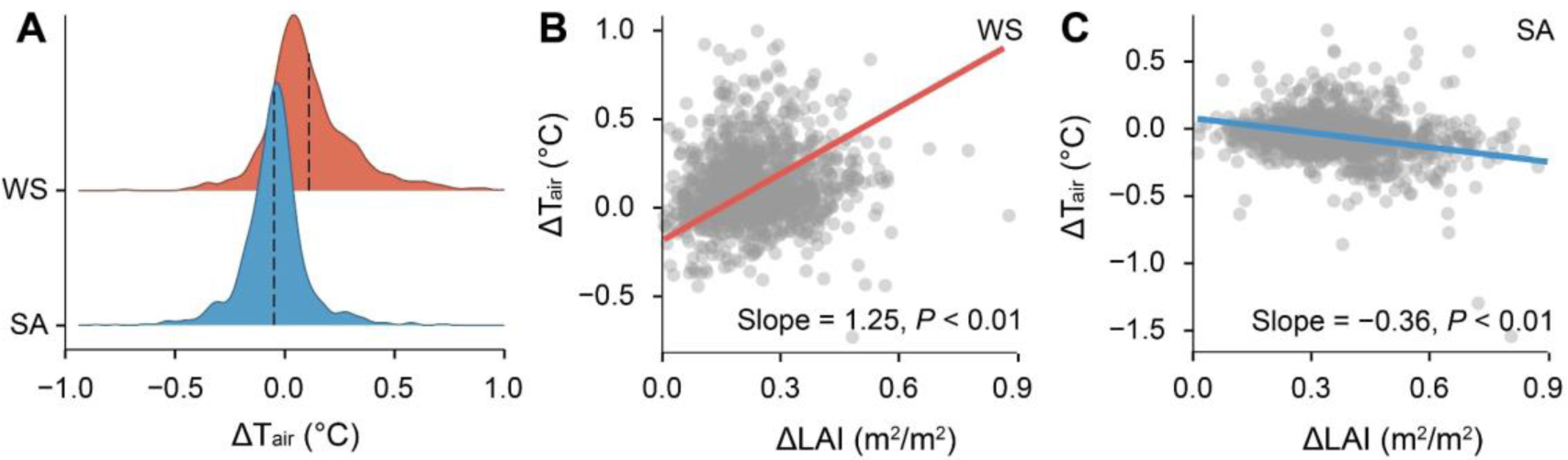
Effect of greening (leaf area index differences, ΔLAI) on temperature gradient (air temperature differences, ΔT_air_) in temperate and boreal forests over the study period 2001-2021. **A–C**, Density plot of ΔT_air_ between WS (winter and spring) and SA (summer and autumn) (**A**), and changes in ΔT_air_ with the increase in ΔLAI in WS (**B**) and in SA (**C**). In **A**, the black dash lines represent mean air temperature gradients for two growing seasons. In **B** and **C**, the circles represent the values of mean ΔLST in WS and SA at each window.

**Fig. S2.**
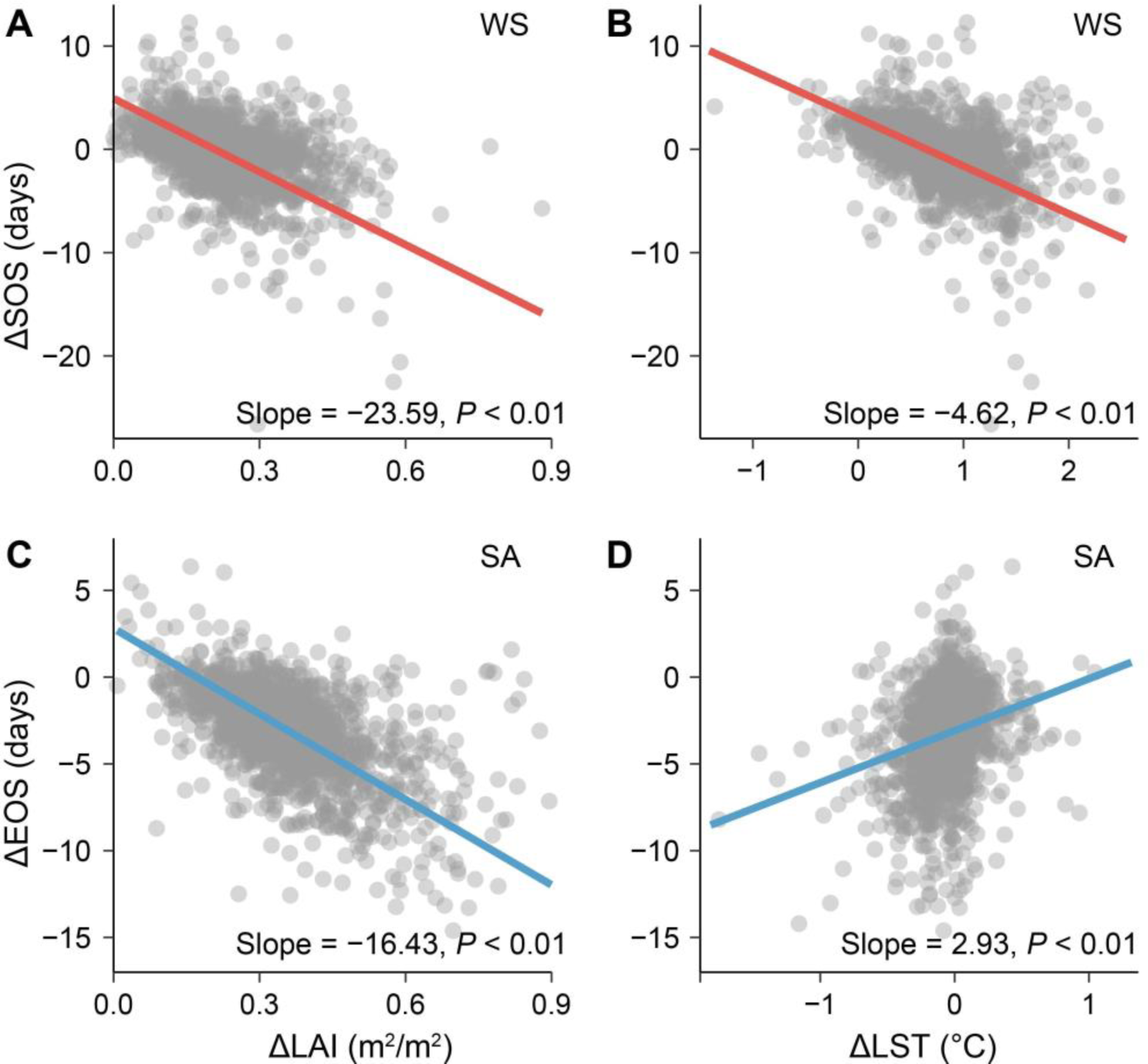
Effects of greening (leaf area index differences, ΔLAI) and greening-induced temperature gradient (land surface temperature differences, ΔLST) for two growing seasons on NIRv-derived phenological differences across different greening gradients in temperate and boreal forests over the study period 2001-2021. **A–D**, Changes in ΔSOS with the increase in ΔLAI (**A**) and ΔLST (**B**) in winter and spring (WS, November to April), changes in ΔEOS with the increase in ΔLAI (**C**) and ΔLST (**D**) in winter and spring (SA, May to October). In **A** to **D**, the circles represent the values of mean ΔSOS in WS and ΔEOS in SA at each window.

**Fig. S3.**
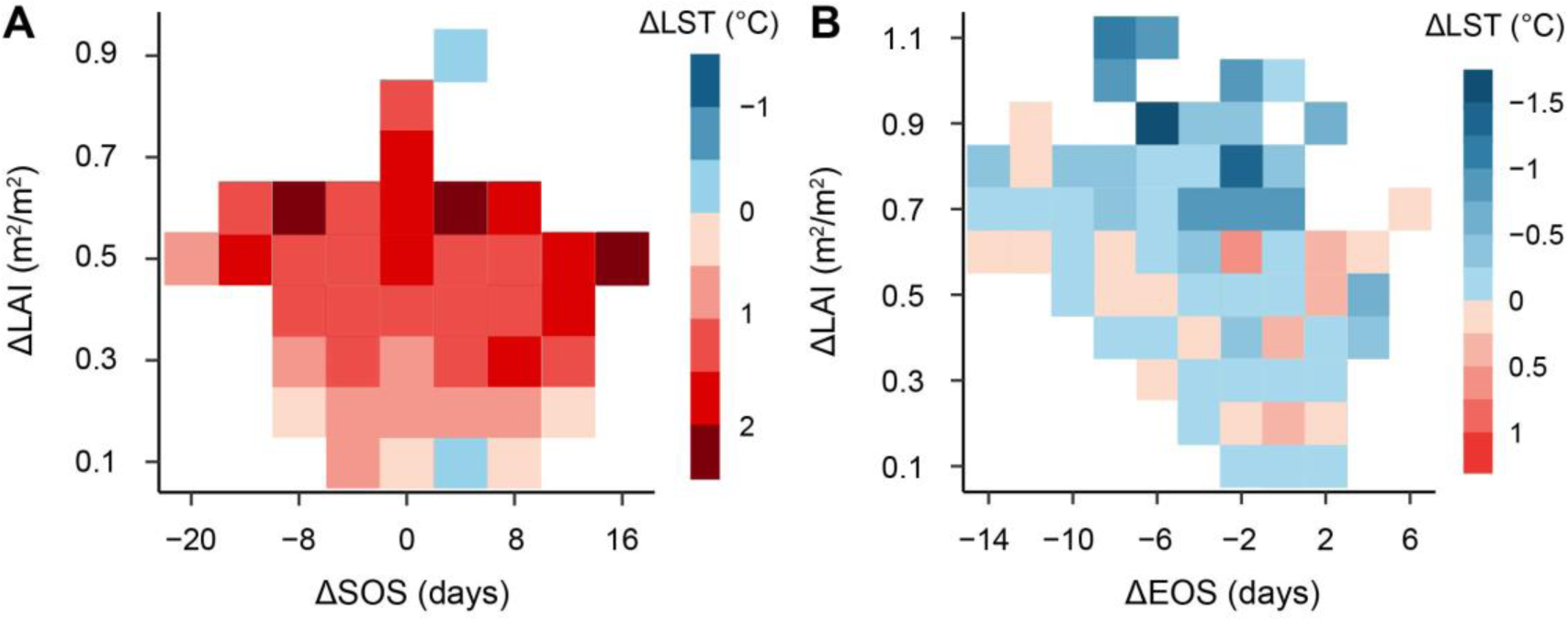
Average metrics for two varying variable gradients. **A–B**, averaged value of ΔLST (**A**) for varying mean ΔLAI and ΔSOS in winter and spring, and averaged value of ΔLST (**B**) for varying mean ΔLAI and ΔEOS in summer and autumn.

**Fig. S4.**
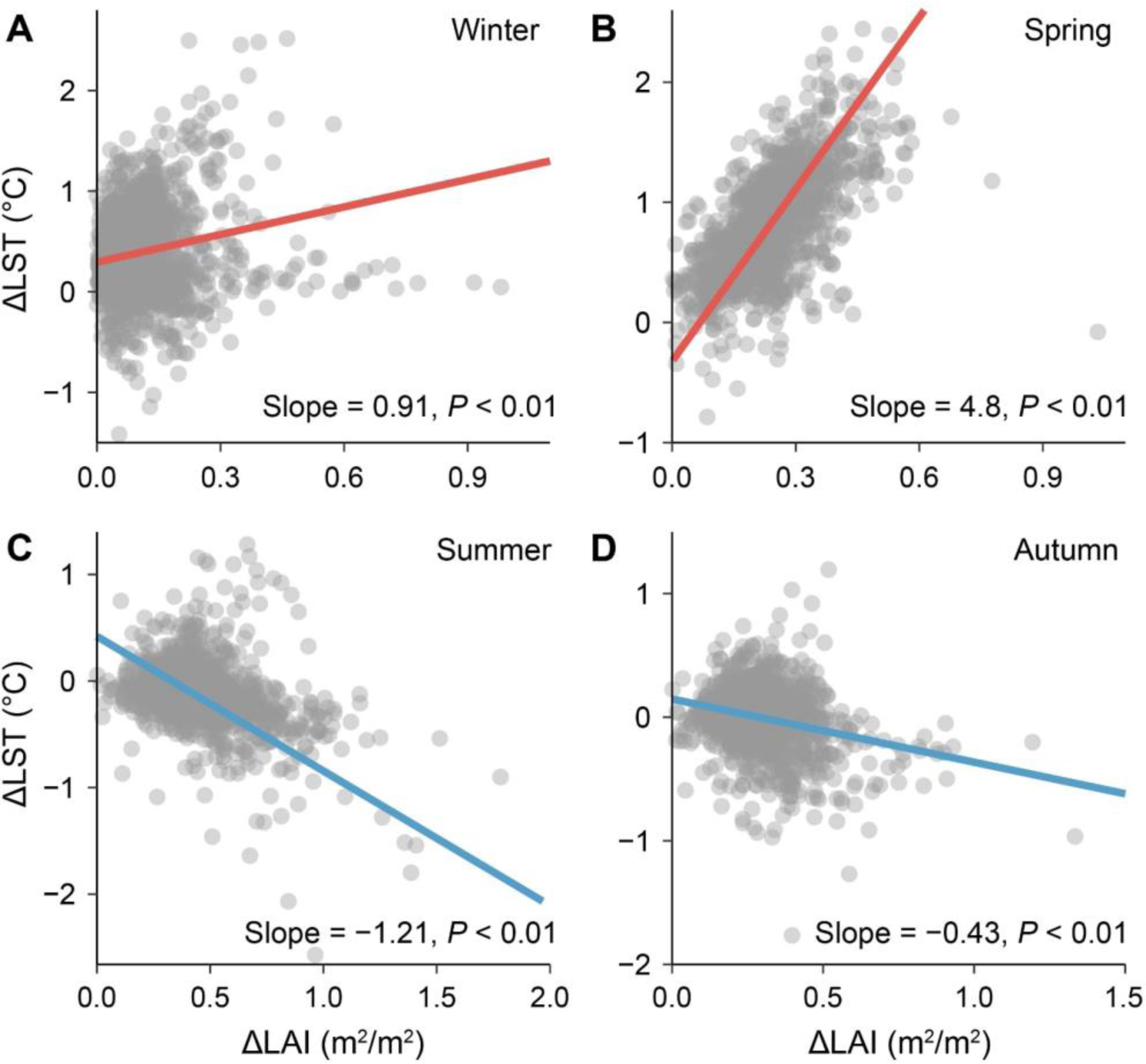
Effect of greening (leaf area index differences, ΔLAI) on temperature gradient (land surface temperature differences, ΔLST) for single seasons in temperate and boreal forests over the study period 2001-2021. **A–D**, Winter (November to January) (**A**), spring (February to April) (**B**), summer (May to July) (**C**), and autumn (August to October) (**D**). In **A** to **D**, the circles represent the values of mean ΔLST for single seasons at each window.

**Fig. S5.**
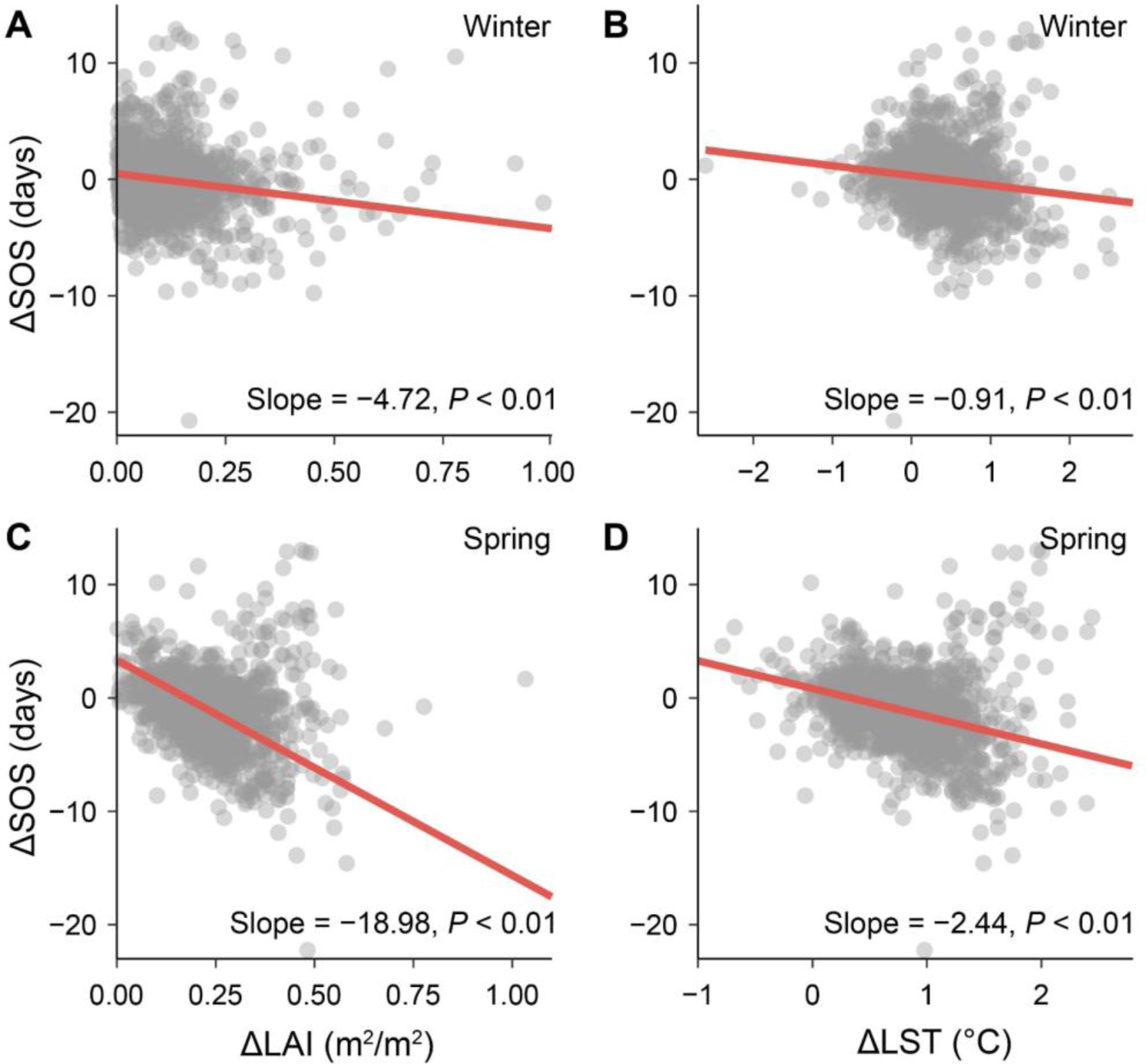
Effects of greening (leaf area index differences, ΔLAI) and greening-induced temperature gradient (land surface temperature differences, ΔLST) for single seasons on the autumn phenology differences (ΔSOS) across different greening gradients in temperate and boreal forests over the study period 2001-2021. **A–C**, Changes in ΔSOS with increased ΔLAI in winter (November to January) (**A**) and spring (February to April) (**C**), and changes in ΔSOS with increased greening-induced ΔLST in winter (**B**) and spring (**D**). In **A** to **D**, the circles represent the values of mean ΔSOS in a single season at each window.

**Fig. S6.**
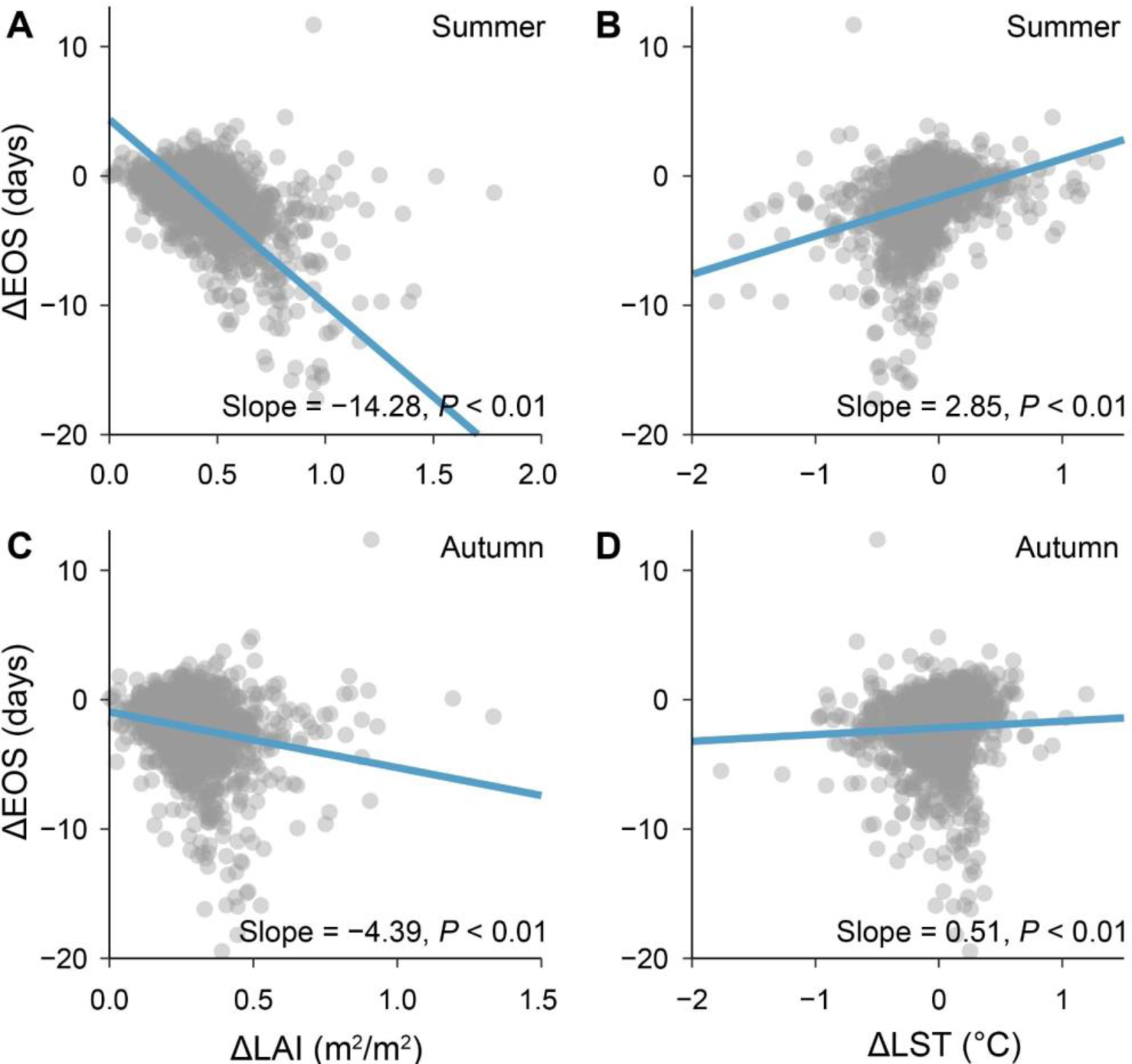
Effect of greening (leaf area index differences, ΔLAI) and greening-induced temperature gradient (land surface temperature differences, ΔLST) for single seasons on spring phenology differences (ΔSOS) across different greening gradients in temperate and boreal forests over the study period 2001-2021. **A–C**, Changes in ΔEOS with increased ΔLAI in summer (May to July) (**A**) and autumn (August to October) (**C**), and changes in ΔEOS with increased greening-induced ΔLST in summer (**B**) and autumn (**D**). In **A** to **D**, the circles represent the values of mean ΔEOS in a single season at each window.

**Fig. S7.**
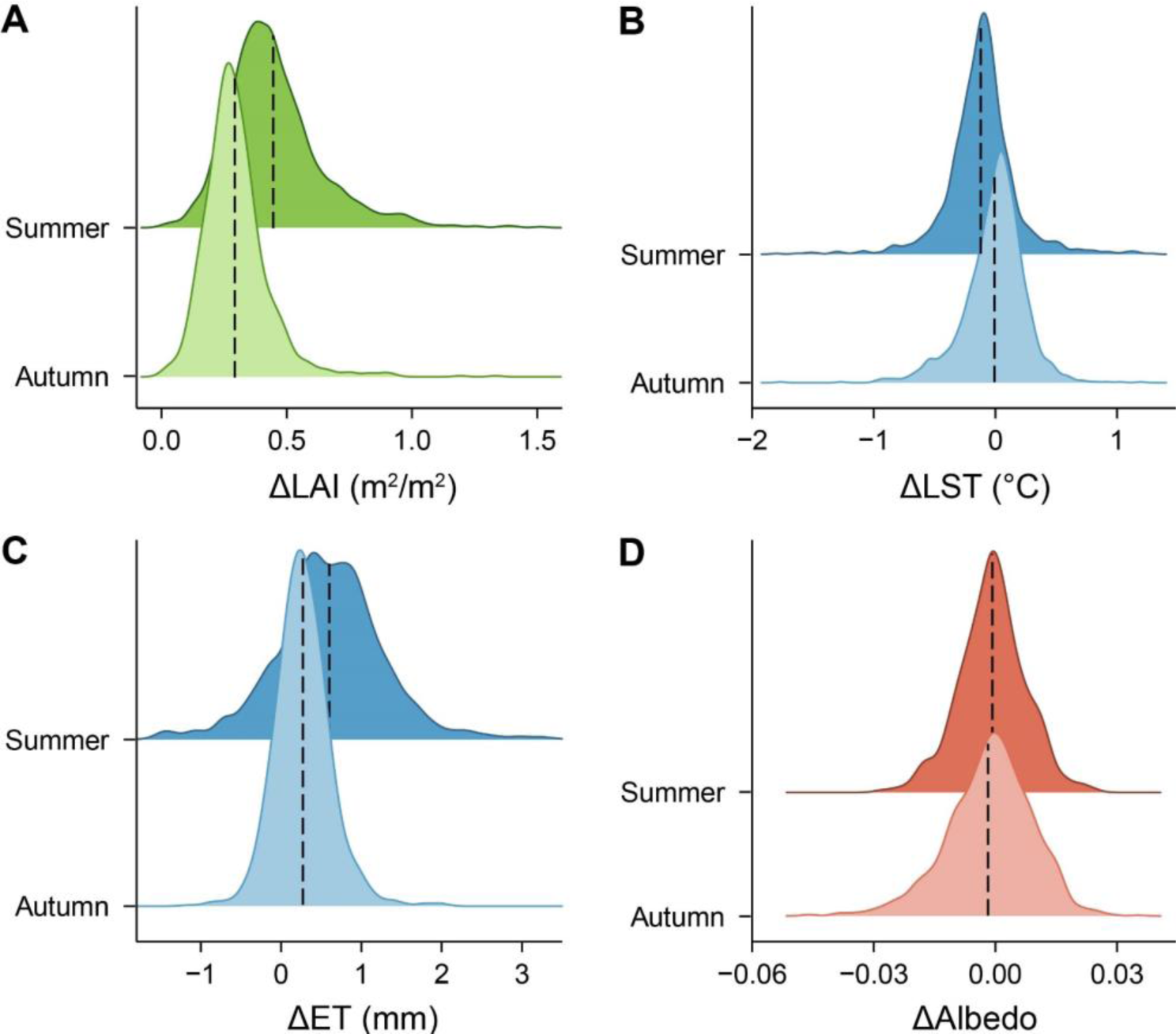
Density plots of mean greening (leaf area index differences, ΔLAI), land surface temperature gradient (ΔLST), evapotranspiration gradient (ΔET), and albedo gradient (ΔAlbedo) for single seasons in temperate and boreal forests during the study period 2001-2021. **A**–**D**, Changes in ΔLAI (**A**) ΔLST (**C**), ΔET (**B**), and ΔAlbedo (**D**) between summer and autumn. In **A** to **D**, the black dash lines represent mean **Δ**LAI, **Δ**LST, ΔET, and ΔAlbedo in summer and in autumn.

**Fig. S8.**
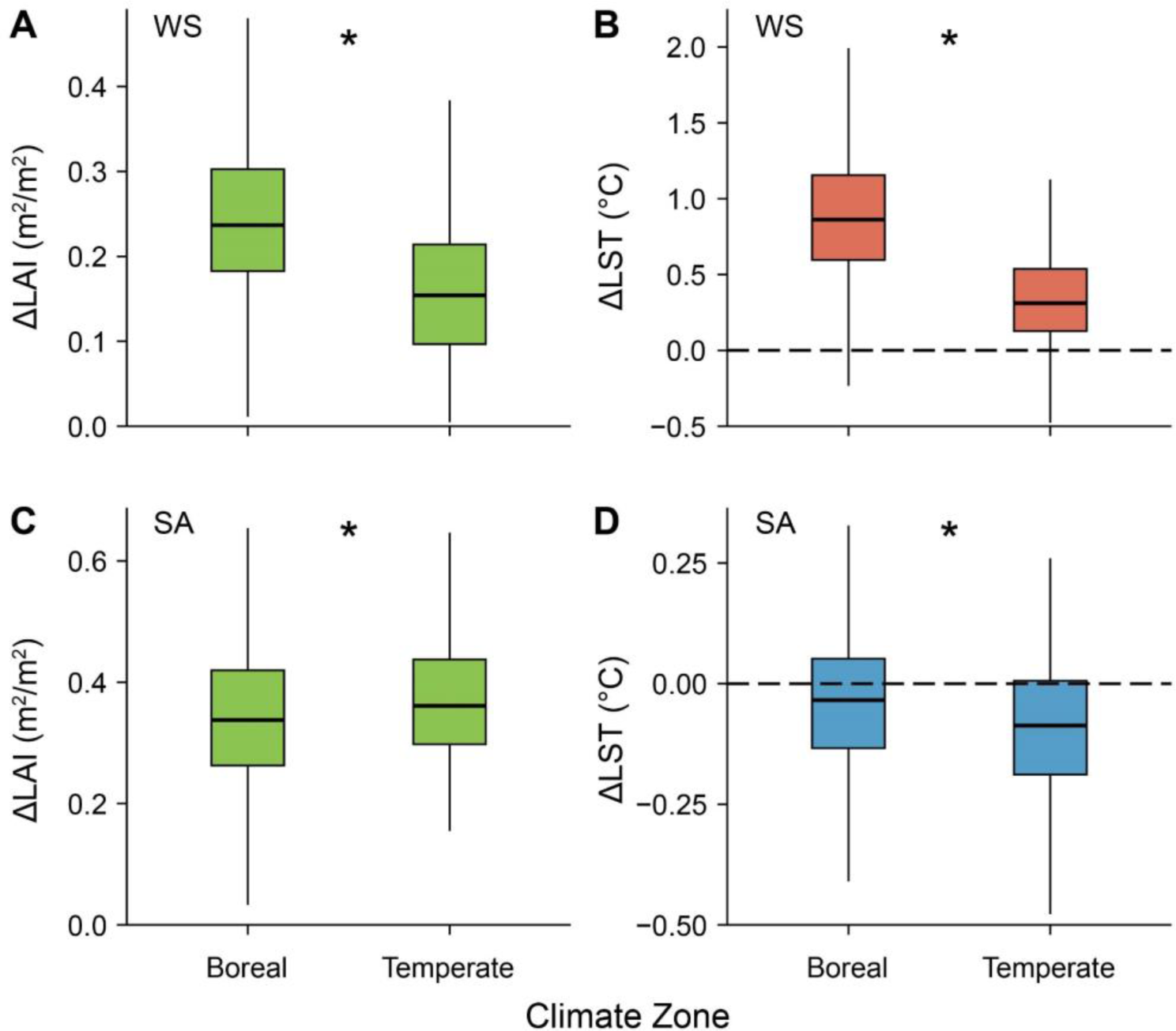
Changes in mean greening (leaf area index differences, ΔLAI) and greening-induced temperature gradient (land surface temperature differences, ΔLST) for two growing seasons between forests with high and low greening gradients in temperate and boreal forests over the study period 2001-2021 across different climate zones. **A–D**, ΔLAI in winter and spring (WS, November to April) (**A**) and in summer and autumn (SA, May to October) (**C**), changes in ΔLST in WS (**B**) and Δ in SA (**D**). In **B** and **D**, the black dash lines represent mean ΔLST in WS and in SA, respectively. In **A** to **D**, the length of each box indicates the interquartile range, the horizontal line inside each box the median, and the bottom and top of the box the first and third quartiles respectively. The asterisk indicates a significant difference in ΔLAI and ΔLST in WS and in SA between boreal and temperate aeras (*P*<0.01).

**Fig. S9.**
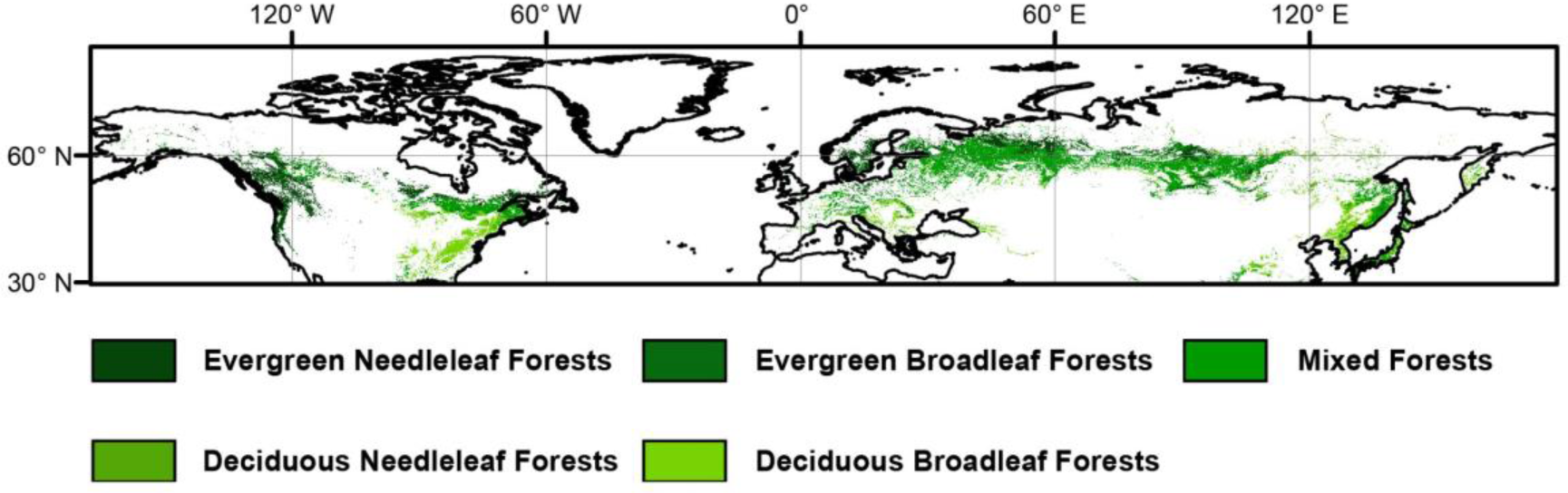
Map of forest types in the Northern Hemisphere derived from MCD12C1 products.

**Fig. S10.**
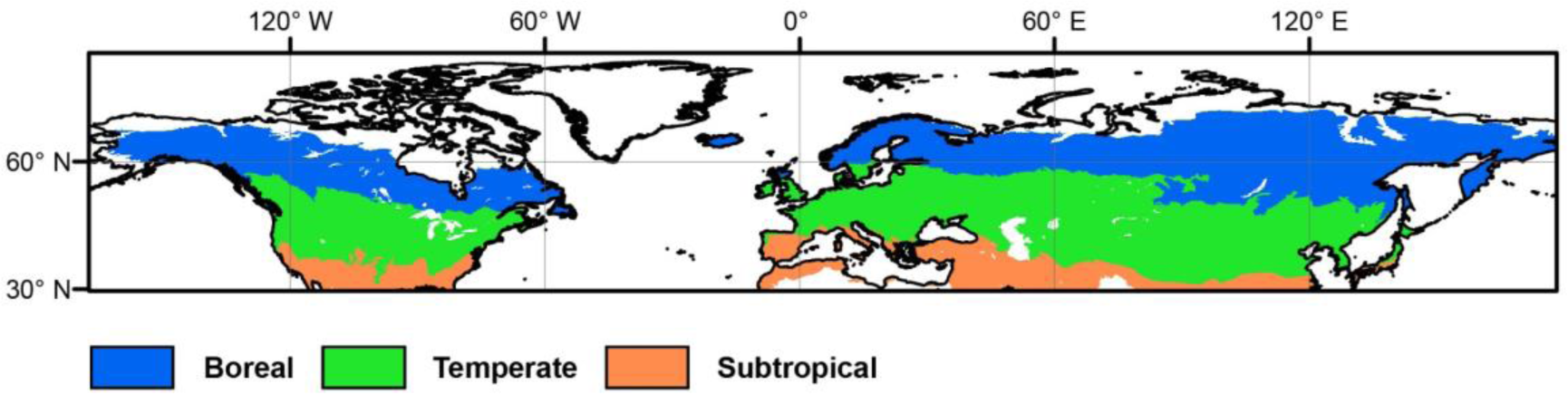
Three major climate zones in the Northern Hemisphere aggregated from global ecological zone (GEZ) map.

**Table S1.**
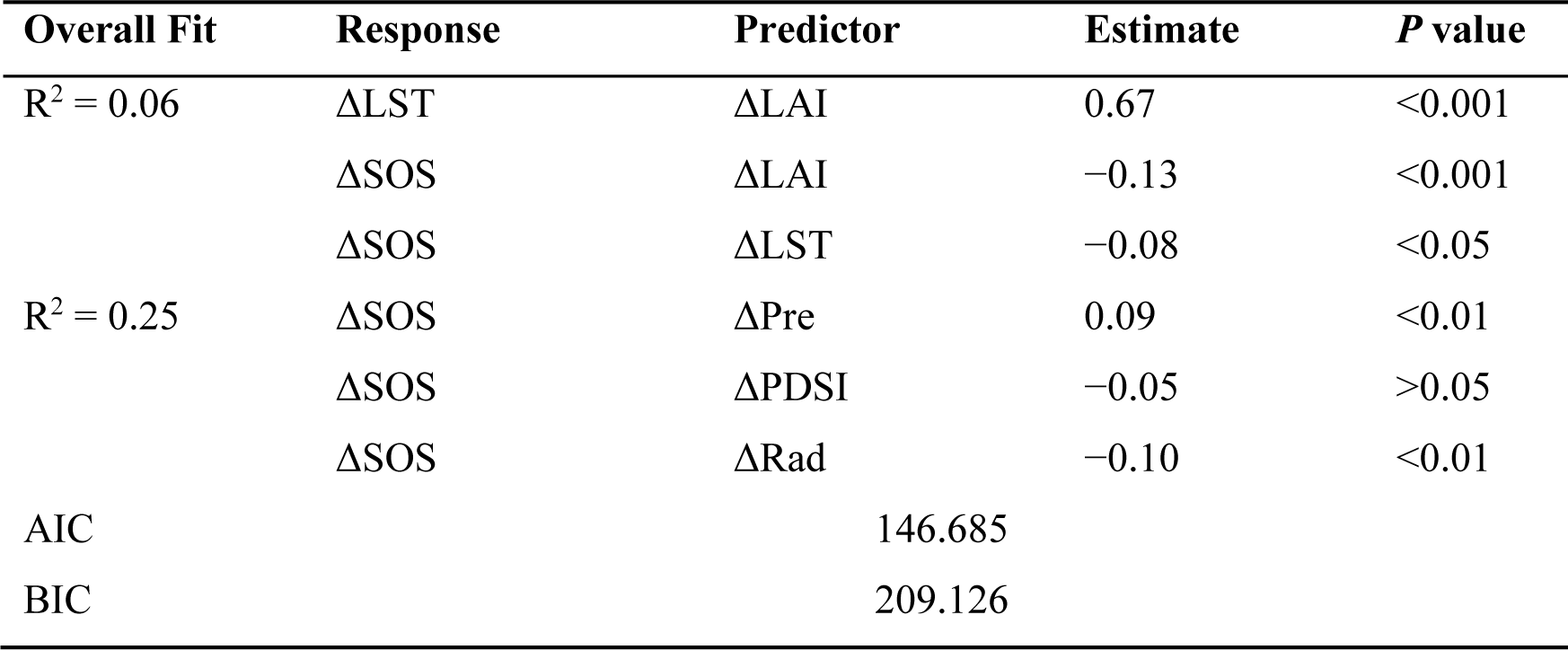
Statistics of the piecewise structural equation model (SEM). SEM was used to explore the direct or indirect effects of greening (ΔLAI), and climate variables gradients on ΔSOS. In the direct-effect model, the ΔLAI, ΔLST, ΔPre, ΔPDSI, and ΔRad in winter and spring (WS, November to April) were assumed to have a direct influence on ΔSOS. In the indirect-effect model, the ΔLAI was assumed to influence ΔSOS by altering ΔLST in WS. We calculated the adjusted coefficients of predictors (R^2^) in each model. The value of standardized direct effect represents the effect of the predictors on the responses. The two-sided test was used to calculate *P* values.

**Table S2.** Statistics of the piecewise structural equation model (SEM). SEM was used to explore the direct or indirect effects of greening (ΔLAI), and climate variables gradients on ΔSOS. In the direct-effect model, the ΔLAI, ΔLST, ΔPre, ΔPDSI, and ΔRad in summer and autumn (SA, May to October) were assumed to have a direct influence on ΔEOS. In the indirect-effect model, the ΔLAI was assumed to influence ΔEOS by altering ΔLST in SA. We calculated the adjusted coefficients of predictors (R^2^) in each model. The value of standardized direct effect represents the effect of the predictors on the responses. The two-sided test was used to calculate *P* values.

| Overall Fit | Response | Predictor | Estimate | $P$ value |
| --- | --- | --- | --- | --- |
| $R^2 = 0.06$ | $\Delta\text{LST}$ | $\Delta\text{LAI}$ | -0.24 | <0.001 |
| | $\Delta\text{EOS}$ | $\Delta\text{LAI}$ | -0.44 | <0.001 |
| | $\Delta\text{EOS}$ | $\Delta\text{LST}$ | 0.19 | <0.001 |
| $R^2 = 0.25$ | $\Delta\text{EOS}$ | $\Delta\text{Pre}$ | 0.08 | <0.01 |
| | $\Delta\text{EOS}$ | $\Delta\text{PDSI}$ | -0.02 | >0.05 |
| | $\Delta\text{EOS}$ | $\Delta\text{Rad}$ | 0.03 | >0.05 |
| AIC |  | 261.128 |  |  |
| BIC |  | 322.927 |  |  |

## Reference

1. I. Chuine, E. G. Beaubien, Phenology is a major determinant of tree species range. Ecol. Lett. 4, 500–510 (2001).

2. R. A. Montgomery, K. E. Rice, A. Stefanski, R. L. Rich, P. B. Reich, Phenological responses of temperate and boreal trees to warming depend on ambient spring temperatures, leaf habit, and geographic range. Proc. Natl. Acad. Sci. 117, 10397–10405 (2020).

3. S. J. Thackeray, et al., Phenological sensitivity to climate across taxa and trophic levels. Nature 535, 241–245 (2016).

4. L. Chen, et al., Leaf senescence exhibits stronger climatic responses during warm than during cold autumns. *Nat*. Clim. Change 10, 777–780 (2020).

5. S. Piao, et al., Leaf onset in the northern hemisphere triggered by daytime temperature. Nat. Commun. 6, 6911 (2015).

6. G. Forzieri, R. Alkama, D. G. Miralles, A. Cescatti, Satellites reveal contrasting responses of regional climate to the widespread greening of Earth. Science 356, 1180–1184 (2017).

7. Z. Zeng, et al., Climate mitigation from vegetation biophysical feedbacks during the past three decades. *Nat*. Clim. Change 7, 432–436 (2017).

8. Z. Zhu, et al., Greening of the Earth and its drivers. *Nat*. Clim. Change 6, 791–795 (2016).

9. Y. Li, et al., Local cooling and warming effects of forests based on satellite observations. Nat. Commun. 6, 6603 (2015).

10. Y. Li, et al., Biophysical impacts of earth greening can substantially mitigate regional land surface temperature warming. Nat. Commun. 14, 121 (2023).

11. X. Lian, et al., Biophysical impacts of northern vegetation changes on seasonal warming patterns. Nat. Commun. 13, 3925 (2022).

12. R. Alkama, A. Cescatti, Biophysical climate impacts of recent changes in global forest cover. Science 351, 600–604 (2016).

13. X. Lee, et al., Observed increase in local cooling effect of deforestation at higher latitudes. Nature 479, 384–387 (2011).

14. S.-S. Peng, et al., Afforestation in China cools local land surface temperature. Proc. Natl. Acad. Sci. 111, 2915–2919 (2014).

15. G. B. Bonan, Forests and Climate Change: Forcings, Feedbacks, and the Climate Benefits of Forests. Science 320, 1444–1449 (2008).

16. X. Lee, et al., Observed increase in local cooling effect of deforestation at higher latitudes. Nature 479, 384–387 (2011).

17. Y. H. Fu, et al., Declining global warming effects on the phenology of spring leaf unfolding. Nature 526, 104–107 (2015).

18. J. Wang, et al., Contrasting temporal variations in responses of leaf unfolding to daytime and nighttime warming. Glob. Change Biol. 27, 5084–5093 (2021).

19. C. Wu, et al., Contrasting responses of autumn-leaf senescence to daytime and night-time warming. *Nat*. Clim. Change 8, 1092–1096 (2018).

20. L. Chen, et al., Long-term changes in the impacts of global warming on leaf phenology of four temperate tree species. Glob. Change Biol. 25, 997–1004 (2019).

21. M. Shen, et al., Strong impacts of daily minimum temperature on the green-up date and summer greenness of the Tibetan Plateau. Glob. Change Biol. 22, 3057–3066 (2016).

22. L. Chen, et al., Immediate and carry-over effects of late-spring frost and growing season drought on forest gross primary productivity capacity in the Northern Hemisphere. Glob. Change Biol. 00, 1–17 (2023).

23. D. Gaumont-Guay, H. A. Margolis, F. J. Bigras, F. Raulier, Characterizing the frost sensitivity of black spruce photosynthesis during cold acclimation. Tree Physiol. 23, 301– 311 (2003).

24. O. Sperling, J. M. Earles, F. Secchi, J. Godfrey, M. A. Zwieniecki, Frost Induces Respiration and Accelerates Carbon Depletion in Trees. PLOS ONE 10, e0144124 (2015).

25. S. Piao, et al., Plant phenology and global climate change: Current progresses and challenges. Glob. Change Biol. 25, 1922–1940 (2019).

26. Y. Vitasse, C. Signarbieux, Y. H. Fu, Global warming leads to more uniform spring phenology across elevations. Proc. Natl. Acad. Sci. 115, 1004–1008 (2018).

27. C. M. Zohner, et al., Late-spring frost risk between 1959 and 2017 decreased in North America but increased in Europe and Asia. Proc. Natl. Acad. Sci. 117, 12192–12200 (2020).

28. H. Zhang, I. Chuine, P. Regnier, P. Ciais, W. Yuan, Deciphering the multiple effects of climate warming on the temporal shift of leaf unfolding. *Nat*. Clim. Change 12, 193–199 (2022).

29. M. Friedl, D. Sulla-Menashe, MODIS/Terra+Aqua Land Cover Type Yearly L3 Global 500m SIN Grid V061 [Data set]. NASA EOSDIS Land Process. DAAC (2022).

30. R. Myneni, Y. Knyazikhin, T. Park, MODIS/Aqua Leaf Area Index/FPAR 8-Day L4 Global 500m SIN Grid V061 [Data set]. NASA EOSDIS Land Process. DAAC (2021).

31. M. Friedl, J. Gray, D. Sulla-Menashe, MODIS/Terra+Aqua Land Cover Dynamics Yearly L3 Global 500m SIN Grid V061 [Data set]. NASA EOSDIS Land Process. DAAC (2022).

32. C. Leys, C. Ley, O. Klein, P. Bernard, L. Licata, Detecting outliers: Do not use standard deviation around the mean, use absolute deviation around the median. J. Exp. Soc. Psychol. 49, 764–766 (2013).

33. J. Zhang, et al., NIRv and SIF better estimate phenology than NDVI and EVI: Effects of spring and autumn phenology on ecosystem production of planted forests. Agric. For. Meteorol. 315, 108819 (2022).

34. C. Schaaf, Z. Wang, MODIS/Terra+Aqua BRDF/Albedo Nadir BRDF Adjusted Ref Daily L3 Global – 500m V061 [Data set]. NASA EOSDIS Land Process. DAAC (2021).

35. G. Badgley, C. B. Field, J. A. Berry, Canopy near-infrared reflectance and terrestrial photosynthesis. Sci. Adv. 3, e1602244 (2017).

36. X. Wang, et al., Validation of MODIS-GPP product at 10 flux sites in northern China. Int. J. Remote Sens. 34, 587–599 (2013).

37. X. Wang, et al., No trends in spring and autumn phenology during the global warming hiatus. Nat. Commun. 10, 2389 (2019).

38. FAO, Global Forest Resources Assessment 2010. FAO For. Pap. 163 (2010).

39. Z. Wan, S. Hook, G. Hulley, MODIS/Terra Land Surface Temperature/Emissivity 8-Day L3 Global 1km SIN Grid V061 [Data set]. NASA EOSDIS Land Process. DAAC (2021).

40. S. Running, Z. Mu, M. Zhao, MOD16A2 MODIS/Terra Net Evapotranspiration 8-Day L4 Global 500m SIN Grid V006. NASA EOSDIS Land Process. DAAC (2017).

41. C. Schaaf, Z. Wang, MODIS/Terra+Aqua BRDF/Albedo Daily L3 Global – 500m V061 [Data set]. NASA EOSDIS Land Process. DAAC (2021).

42. J. T. Abatzoglou, S. Z. Dobrowski, S. A. Parks, K. C. Hegewisch, TerraClimate, a high-resolution global dataset of monthly climate and climatic water balance from 1958–2015. Sci. Data 5, 170191 (2018).

43. D. B. Gesch, K. L. Verdin, S. K. Greenlee, New land surface digital elevation model covers the Earth. Eos Trans. Am. Geophys. Union 80, 69–70 (1999).

44. C. A. Gunderson, et al., Forest phenology and a warmer climate – growing season extension in relation to climatic provenance. Glob. Change Biol. 18, 2008–2025 (2012).

45. Q. Liu, et al., Delayed autumn phenology in the Northern Hemisphere is related to change in both climate and spring phenology. Glob. Change Biol. 22, 3702–3711 (2016).

46. R. M. Marchin, C. F. Salk, W. A. Hoffmann, R. R. Dunn, Temperature alone does not explain phenological variation of diverse temperate plants under experimental warming. Glob. Change Biol. 21, 3138–3151 (2015).

47. J. S. Lefcheck, piecewiseSEM: Piecewise structural equation modelling in r for ecology, evolution, and systematics. Methods Ecol. Evol. 7, 573–579 (2016).

48. R Core Team, R: A language and environment for statistical computing. *R Found*. Stat. Comput. (2021).

49. J. Elith, J. R. Leathwick, T. Hastie, A working guide to boosted regression trees. J. Anim. Ecol. 77, 802–813 (2008).

50. Ridgeway G, Ridgeway M.G, The gbm package. R Found. Stat. Comput., 5(3) (2004).

51. N. Gorelick, et al., Google Earth Engine: Planetary-scale geospatial analysis for everyone. Remote Sens. Environ. 202, 18–27 (2017).

